# A Cryptic Binding Pocket Regulates the Metal-Dependent Activity of Cas9

**DOI:** 10.1101/2025.08.25.672025

**Authors:** Mohd Ahsan, Aakash Saha, Delisa Ramos, Isabel Strohkendl, Alexa L. Knight, Erin Skeens, George P. Lisi, David W. Taylor, Giulia Palermo

## Abstract

Cas9 is a metal-dependent nuclease that has revolutionized gene editing across diverse cells and organisms exhibiting varying ion uptake, metabolism, and concentrations. However, how divalent metals impact its catalytic function, and consequently its editing efficiency in different cells, remains unclear. Here, extensive molecular simulations, Markov State Models, biochemical and NMR experiments, demonstrate that divalent metals – Mg^2+^, Ca^2+^, and Co^2+^ – promote activation of the catalytic HNH domain by binding within a dynamically forming divalent metal binding pocket (DBP) at the HNH-RuvC interface. Mutations in DBP residues disrupt HNH activation and impair the coupled catalytic activity of both nucleases, identifying this cryptic DBP as a key regulator of Cas9’s metal-dependent activity. The ionic strength thereby promotes Cas9’s conformational activation, while its catalytic activity is metal-specific. These findings are critical to improving the metal-dependent function of Cas9 and its use for genome editing in different cells and organisms.

---

CRISPR-Cas9 (Clustered Regularly Interspaced Short Palindromic Repeats and associated protein Cas9) has revolutionized life sciences by introducing a facile genome editing technology^1,2^. At its core, Cas9 is a metal-dependent nuclease enabling precise genetic modifications across a wide range of cells and organisms with diverse ion uptake, metabolism, and intracellular metal concentrations^3^. A key distinction between bacterial and mammalian cells is the availability of free divalent ions, which can influence protein-nucleic acid interactions and the specificity of Cas effectors^4–7^. However, the extent to which divalent metals regulate the catalytic function of Cas9, and consequently its genome editing capability in different cellular environments, remains unclear. Understanding the metal-dependent function of Cas9 is therefore important to fully exploit its function across different biological applications, aiding more precise and versatile genome editing technologies.

The *S. pyogenes* Cas9 (SpCas9) is an RNA-guided nuclease that utilizes the sequence specificity of its guide RNA to target and cleave complementary DNA sequences in the presence of metal ions^8^. Double-stranded DNA cleavage is carried out by two catalytic domains, HNH and RuvC (Fig. 1a), which cleave the DNA target strand (TS, complementary to the guide RNA) and the non-target strand (NTS), respectively^3,9^. The HNH domain exhibits considerable conformational flexibility and structural rearrangements, in contrast to the RuvC domain, which maintains a stable structure^10^. Upon DNA binding, the HNH nuclease transitions from an inactive state, in which its catalytic triad (comprising H840, N863, D839) is oriented away from the cleavage site on the DNA TS^11,12^, to a pre-active state, where H840 shifts closer to the scissile phosphate but remains ∼16 Å away (Fig. 1a, left)^13^. While this transition can occur in the absence of metal ions, the final shift from the pre-active to the active state, where H840 docks at the cleavage site (Fig. 1a, right)^14–16^, is dependent on Mg^2+^ ions^17–19^. Single-molecule studies have shown that Mg^2+^ concentrations between 5 and 10 mM (i.e., high concentrations) are required for HNH to transition into its active state^18^. At low Mg^2+^ concentrations (10 μM), this conformational shift is hindered, preventing HNH activation for DNA cleavage. These studies also revealed that other divalent cations, such as Ca^2+^ and Co^2+^, can promote this conformational activation when present at high concentrations. These findings suggest that high concentrations of divalent cations may promote the docking of HNH at the DNA target strand for cleavage. However, the impact of ionic strength on this critical conformational change, which is essential for Cas9 function, remains unclear. Additionally, how different divalent metal ions – such as Mg^2+^, Ca^2+^, and Co^2+^ – alter the chemical mechanism of DNA cleavage, either facilitating or inhibiting catalysis^3^, remains an open question. Addressing these questions is key to understanding how Cas9 function is altered in cells with varying ion uptake, metabolism, and intracellular concentrations, as well as for optimizing Cas9-mediated genome editing across diverse biological contexts, including plants and mammalian cells.

**Figure 1.**
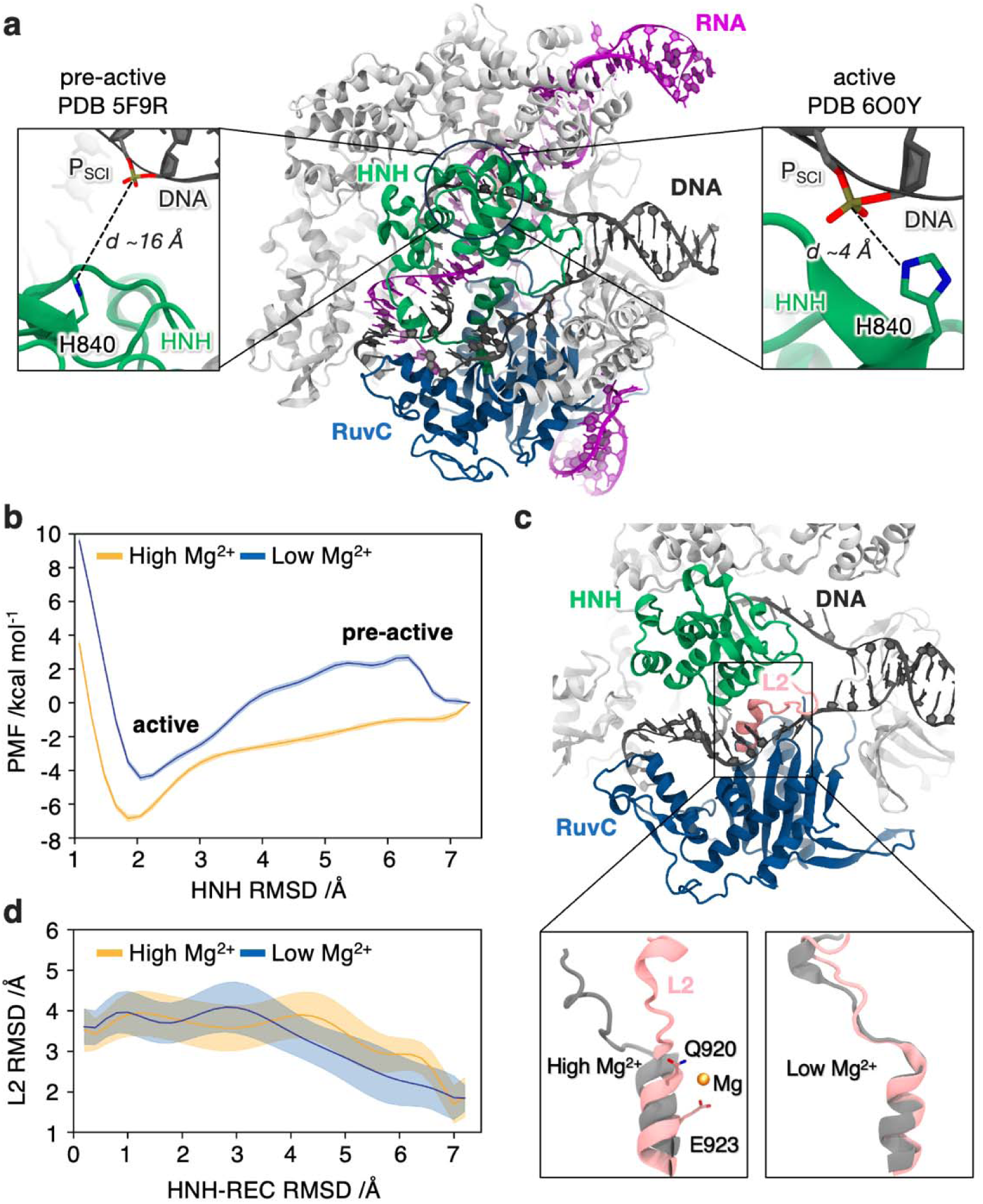
Cas9 activation under varying concentrations of Mg^2+^ ions. **a.** Structure of *S. pyogenes* Cas9 bound to RNA (magenta) and DNA (black), highlighting the HNH (green) and RuvC (blue) catalytic domains^14^. Close-up views show the HNH nuclease in its pre-active state (PDB 5F9R^13^, left), where the catalytic H840 is ∼16 Å away from the scissile phosphate (P_SCI_) on the DNA target strand, and in its active conformation (PDB 6O0Y^14^, right), where HNH is docked at the cleavage site. **b.** Free energy profiles depicting the conformational change of HNH from its pre-active to active state at high (10 mM, yellow) and low (10 μM, blue) concentration of Mg^2+^. The Potential of Mean Force (PMF, in kcal/mol) was computed along the root-mean-square deviation (RMSD) of Cα atoms in the HNH domain and portions of the recognition lobe (REC) relative to both states. Error estimation was performed using Monte Carlo bootstrap analysis of free energy simulations (see Supplementary Methods). **c.** Rearrangements of the L2 region (residues 907–925) in response to Mg^2+^ concentration. At high Mg^2+^ levels (left), residues Q920 and E923 form interactions with the ion, which are lost under low Mg^2+^ conditions (right; Supplementary Fig. 3). **d.** RMSD of the L2 region along the free energy profiles reveals that as HNH moves toward the active state (i.e., in the transition region from RC ∼7 to ∼4 Å), conformational changes in L2 are more pronounced at high Mg^2+^ concentration. Errors are reported as the standard deviation for each free energy simulation window.

Here, extensive molecular simulations, including free energy methods and Markov State Models, integrated with biochemical experiments and solution NMR, characterize the thermodynamics and kinetics of the HNH activation under varying concentrations of metals. We reveal that divalent metals facilitate HNH activation and the coordinated function of both nucleases by binding within a divalent metal binding pocket (DBP), which dynamically forms at the HNH-RuvC interfaces. Mutations in this cryptic DBP disrupt HNH dynamics and impair the coupled activity of both nucleases, underscoring its regulatory role in Cas9’s metal-dependent function. Quantum-classical simulations further elucidate the metal-specific nature of Cas9’s catalytic activity showing that while Mg^2+^ and Ca^2+^ support DNA cleavage, Co^2+^ disrupts enzymatic function, despite ionic strength promoting Cas9 conformational activation regardless of ion type. These findings advance our understanding of Cas9’s metal-dependent mechanism and offer valuable insights for optimizing genome editing applications.

## Results

### Magnesium-Dependent Transition Pathways of the HNH Domain

The transition of the HNH domain from the pre-active (PDB 5F9R)^13^ to the active (PDB 6O0Y)^14^ state (Fig. 1a) under varying Mg^2+^ concentrations was investigated using a combination of free energy simulations and Markov State Model (MSM) analysis^20,21^, generating ∼12 μs total sampling across systems. The Umbrella Sampling (US) method^22^ was employed to examine the effects of high (10 mM) and low (10 μM) Mg^2+^ concentrations on the energetics of the HNH transition, while a MSM approach provided the transition kinetics upon additional unbiased simulations (see Supplementary Methods). US simulations were performed using the difference in root-mean-square deviation (RMSD) of the Cα atoms in the HNH domain and portions of the recognition lobe relative to the pre-active and active states as the reaction coordinate (RC, Supplementary Fig. 2).

At high Mg^2+^ concentration, the transition from the pre-active to the active state proceeds barrierless, progressing toward the energetic minimum corresponding to the active state (Fig.1b). At low Mg^2+^ concentration, a ∼2.5 kcal/mol energetic barrier indicates a less favourable transition. The Potential of Mean Force (PMF) profile further indicates that the active state is less thermodynamically stable under low Mg^2+^ conditions, underscoring the role of Mg^2+^ in stabilizing the active state. The simulations also uncover structural rearrangements of the L2 region (residues 907–925), which bridges the HNH and RuvC domains (Fig. 1c)^13^. Indeed, as the HNH domain transitions toward the active state (from RC ∼7 to ∼4 Å), conformational changes in L2 are more pronounced at high Mg^2+^ levels (Fig. 1d) and are linked to the coordination of Mg^2+^ ions by residues Q920 and E923 (Fig. 1c, Supplementary Fig. 3).

To gain insights into the transition kinetics, a MSM was employed^20,21^. This approach describes the possible transitions between conformational states and their intermediates, along with the likelihood of these transitions over time. To construct our MSM, additional unbiased MD simulations were performed along the HNH conformational change, yielding ∼3.6 μs of sampling for each Mg^2+^ concentration examined (see Supplementary Methods). The HNH conformational change was described using the Cα–Cα distances between S355 and S867, and between S867 and N1054, which served as independent components (ICs) for constructing our MSM. These distances, previously utilized in Förster Resonance Energy Transfer (FRET) experiments^23^, effectively represent the conformational activation of HNH^10^, and delineate the differences between the investigated states (Supplementary Figs. 4, 5). Our MSM enabled a detailed characterization of the transition kinetics from the pre-active (State A) to the active state (State B, Fig. 2a). At both high and low Mg^2+^ levels, the transition is mediated by two intermediate metastable states (state 1 and state 2, Fig. 2b, Supplementary Figs. 6-8). At high Mg^2+^ concentration, the flux through the slowest transition step is twice as fast, relative to low Mg^2+^ concentration, reinforcing the critical role of Mg^2+^ in driving this transition.

**Figure 2.**
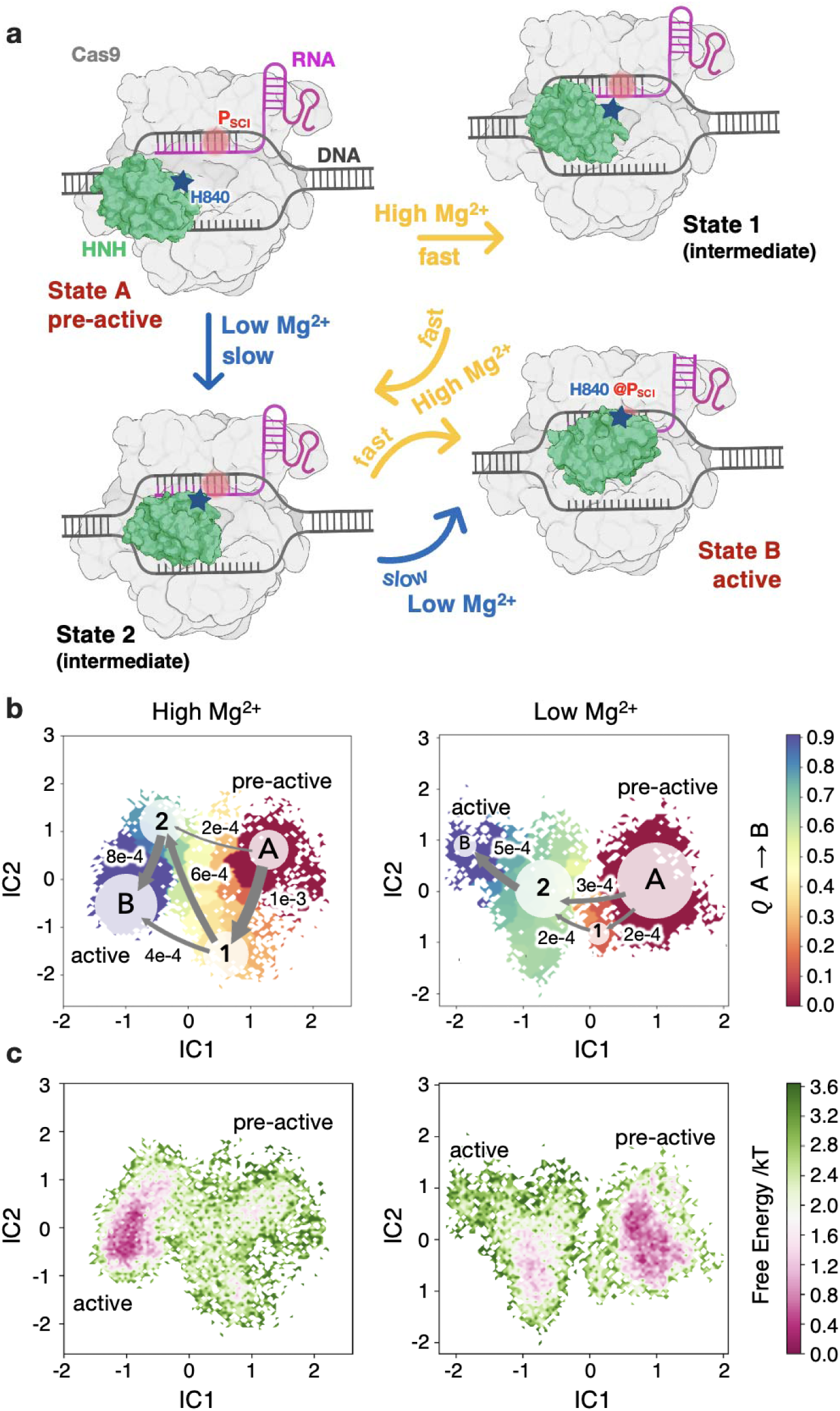
Kinetics of the HNH conformational change. **a.** Schematic representation of the transition kinetics of the HNH domain from the pre-active (State A) to the active (State B) conformation via intermediate states (viz., States 1 and 2). The transition occurs approximately twice as fast (see panel **b**) at high Mg^2+^ concentration (10 mM) compared to low Mg^2+^ levels (10 μM). **b.** Markov State Model (MSM) analysis depicting transitions between states under high (left) and low (right) Mg^2+^ conditions. Distances between residue pairs S355–S867 and S867–N1054 were used as independent components (IC1 and IC2) for time-lagged independent component analysis (TICA)^24^, presented as scatter TICA plots color-coded by the committor probability (*Q*; scale on the right). The *Q* values ∼0.5 indicate that the system at the transition state, traversing two metastable states (State 1 and State 2) *en route* from the pre-active to the active state. Transition flux values show that the conformational change proceeds approximately twice as fast at high Mg^2+^ concentration. **c.** TICA plots color-coded according to the free energy landscape, obtained by reweighting the Boltzmann distribution using stationary probabilities obtained from MSM at high (left) and low (right) Mg^2+^ concentrations. At high Mg^2+^, the active conformation is more frequently sampled and energetically favoured. In contrast, under low Mg^2+^ conditions, the system predominantly resides in the pre-active state.

Analysis of the L2 region revealed marked conformational changes at high Mg^2+^ levels, with Q920 and E923 interacting with Mg^2+^ ions (Supplementary Fig. 9). The free energy landscapes from MSM analysis also show that at high Mg^2+^ levels, the active state (State B) is both energetically favoured and more frequently populated, indicating an efficient transition (Fig. 2c).

At low Mg^2+^ concentration, the system is often trapped in intermediate metastable states, resulting in an incomplete transition to the active state. These findings demonstrate that elevated Mg^2+^ levels not only promote an effective transition to the active state but also stabilizes it.

### Divalent Ions Drive the HNH Transition Through a Cryptic Binding Pocket

To further investigate the influence of divalent metals on the dynamics of HNH, all-atom MD simulations were performed at high (10 mM) and low (10 μM) concentrations of Mg^2+^, Ca^2+^, and Co^2+^, totalling ∼36 μs. These simulations were initiated from the pre-active state of Cas9, providing a view of how varying ionic environments affect key molecular interactions. As a result, Mg^2+^ ions were observed to interact at two primary sites: between the HNH domain and the P_SCI_ and at the junction between HNH and RuvC (Fig. 3a). Under high Mg^2+^ concentrations, HNH displayed a more pronounced movement toward the scissile phosphate (P_SCI_) on the DNA TS, evidenced by the reduction of key distances bridging the HNH catalytic core and the TS (Fig. 3b, Supplementary Fig. 10). This observation is corroborated by the analysis of the backbone RMSD, highlighting that elevated Mg^2+^ concentrations enhance the dynamics of HNH while stabilizing RuvC (Supplementary Fig. 11). High Mg^2+^ concentrations also increased the frequency of bridging interactions between the ion, HNH (through the coordination of D850) and P_SCI_ (Fig. 3c). Similar effects were observed with Ca^2+^ and Co^2+^ (Supplementary Fig. 12), supporting the role of divalent ion binding at this site in facilitating the conformational transition. The second divalent ion binding site emerges as a pocket during molecular simulations, positioned between the HNH-RuvC interface and involving residues from the L2 region, the RuvC domain, and the NTS (Fig. 3a). This Divalent ion Binding Pocket (DBP) recruits Mg^2+^ ions at high concentrations, forming interactions that bridge the L2 region with the RuvC domain and the NTS (Fig. 3d, Supplementary Fig. 13). Simulations employing different Mg^2+^ force-field parameters confirmed that these bridging interactions are more pronounced at high Mg^2+^ concentrations (Supplementary Fig. 14). Their formation of at low Mg^2+^ concentration is hindered by shorter residence times, compared to high Mg^2+^ levels (Fig. 3e). Similar results were observed in the presence of Ca^2+^, and Co^2+^ (Supplementary Fig. 15), indicating that high concentrations of divalent ions – regardless of the specific ion – facilitate the formation of bridging interactions within the DBP. These findings align with the ability of MD simulations to reveal the formation of hidden cavities, *viz.,* “cryptic pockets”, which may not be evident in experimental structures but can provide critical insights into protein function^25–28^. The observed interactions also corroborate the rearrangement of the L2 region, observed during the transition from the pre-active to active state (Fig. 1d). This reorganization, along with the bridging interactions within the cryptic DBP, promotes the activation of the HNH domain. To validate these computational insights, biochemical experiments were conducted. Mutations (mutants M1: D54A/S55A, E57A, M2: Q920A/E923A, M3: T13A/N14A) were introduced into residues lining the DBP, and the resulting plasmid DNA cleavage activity was compared to that of wild-type Cas9. Loss of supercoiled plasmid substrate and appearance of linear product and nicked intermediate were fit to single and double exponential curves, respectively, to determine the apparent rates of the first and second strand cleavage events. All mutants exhibited reduced catalytic efficiency (Fig. 3f, Supplementary Fig. 16).

**Figure 3:**
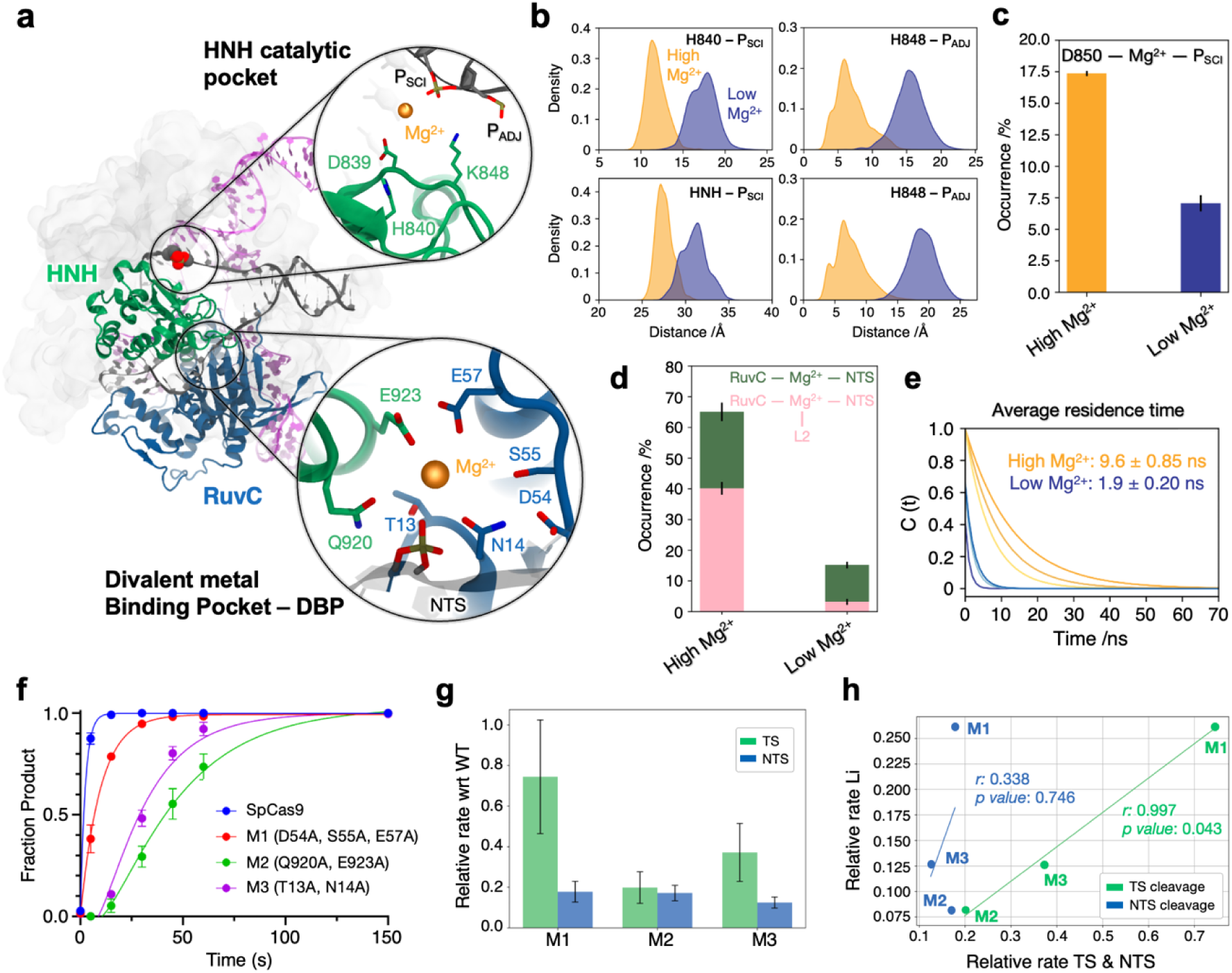
Divalent Ion Binding Sites in Cas9. **a.** Overview of Cas9 highlighting two distinct divalent ion binding sites. The HNH catalytic pocket (close-up view, top) features a divalent ion (Mg^2+^) between the catalytic core and the scissile phosphate (P_SCI_) on the DNA target strand (TS). A cryptic divalent metal binding pocket (DBP, close-up view, bottom) emerges during molecular simulations between the HNH and RuvC domains. This pocket comprises residues of RuvC, the L2 region, and the DNA non-target strand (NTS), with the Mg^2+^ ion bridging interactions among these regions. **b.** Probability distribution of key distances bridging the HNH catalytic core and the TS at high (10 mM, yellow) and low (10 μM, blue) Mg^2+^ concentrations ([Mg^2+^]). Measured distances include those between residues H840 and K848 (CC atoms) with P_SCI_ and the adjacent phosphate (P_ADJ_), and the distance between the centre of mass (COM) of HNH and P_SCI_. **c-d.** Percentage occurrence (with standard error of the mean) of bridging interactions established by Mg^2+^ in the HNH catalytic site (**c**) and within the DBP (**d**) at high and low [Mg^2+^]. In the active site, bridging interactions involve Mg^2+^, D850 and P_SCI_. Within the DBP, interactions are categorized as follows: Mg^2+^ binding the RuvC domain and the NTS (green), and Mg^2+^ bridging RuvC, the NTS and L2 region (pink). **d.** Average residence time of Mg^2+^ ions calculated from the time correlation function (*C_t_*). Data in panels **b-e** are computed considering the overall ensemble, with a comprehensive analysis in Supplementary Figs. 10-15. f. Cleavage activity over time for plasmid DNA by wild-type Cas9 and DBP mutants (see Supplementary Fig. 16 for details). Data points represent the mean of three independent replicates ± standard error of the mean, fitted to a double-exponential curve. g. Calculated rates of TS (green) and NTS (blue) cleavage relative to wild-type Cas9 (*k_obs_*, Supplementary Fig. 17). Rates were determined using fluorescently labelled oligoduplex substrates under saturating conditions to measure TS and NTS cleavage. Error bars represent the standard error of the fit. h. Correlation between the relative rate of appearance of Linear (product) species and the relative rate cleavage activity for both TS (green) and NTS (blue) cleavage (details in Supplementary Fig. 17) with respect to wild-type SpCas9.

Mutant M1 showed an equal reduction in the loss of supercoiled substrate and formation of linear product. Mutants M2 and M3, in addition to dramatically slower plasmid cleavage, also produced a prominent nicked intermediate species (Supplementary Fig. 16), suggesting that the sets of residues mutated within the DBP differ in magnitudes of effect on the RuvC and HNH domains. These observations indicate that the mutations impair cleavage mediated by the HNH and RuvC nuclease domains and suggest a disruption in the communication between them^23,29,30^. Additionally, we performed reactions with fluorescently labelled oligoduplex substrates, enabling separate measurements of DNA TS and NTS cleavage to quantify penalties to the HNH and the RuvC domain, respectively (Supplementary Fig. 17). RuvC NTS cleavage (*k*_obs_) was uniformly reduced by ∼80-85% across all mutants, while HNH TS cleavage (*k*_obs_) was reduced by ∼25%, ∼80%, and ∼63% for mutants M1, M2, and M3, respectively (Fig. 3g). When compared to the formation of linearized product within the plasmid cleavage assay, there is a strong correlation with the relative rate of TS cleavage (Fig. 3h, r=0.998), indicating that the different behaviors of the DBP mutants are due to the penalties on HNH cleavage. The observation that the M2 mutant, which includes alanine mutations of the L2 residues Q920 and E923, demonstrates the strongest reduction in HNH cleavage corroborates the role of the L2 region in HNH activation^15^. These findings align with biochemical and biophysical studies showing that the HNH and RuvC domains operate in a coupled catalytic manner^13,23,29–31^, relying on dynamic crosstalk essential for Cas9 activity.

Collectively, these findings suggest that disruptions of the DBP differentially modulate HNH catalysis depending on the mutation site, while the communication between the HNH and RuvC domains remains impaired across all mutants, leading to a comparable reduction in RuvC catalysis. Solution NMR experiments were performed to probe the effect of Mg^2+^ on an isolated HNH-L2 construct. A distinct structural state was observed in the presence of Mg^2+^ (Supplementary Fig. 18), consistent with computational findings (Fig. 2). The catalytic H840 experiences the largest structural perturbation, with the residues proximal to H840 and Q920 in L2 also exhibiting significant structural changes. Fast timescale (ps–ns) dynamics are largely maintained with Mg^2+^, while μs–ms motions in L2 are attenuated (Supplementary Fig. 19), which may contribute to the stabilization of the active state observed through MD simulations at high Mg^2+^ concentration (Fig. 3c). Solution NMR thereby indicate that Mg^2+^ ions alter structure and dynamics of HNH-L2, in line with computational studies. Collectively, computational findings, consistent by biochemical and NMR experiments, demonstrate that a well-structured DBP enhances coordination between the HNH and RuvC domains of Cas9, facilitating the transition of HNH to its active state.

### Impact of Divalent Metal Ions on Target Strand Cleavage

To investigate how different divalent metal ions affect TS cleavage, the active state of Cas9 ^14^ was simulated for ∼9 μs in the in the presence of Mg^2+^, Ca^2+^, and Co^2+^, revealing their impact on the catalytic site’s structure and dynamics. The 2D-RMSD plots of the active site show that Ca^2+^ preserves a structure similar to that stabilized by Mg^2+^, whereas Co^2+^ induces pronounced structural deviations (Fig. 4a, Supplementary Fig. 20). To substantiate these observations, quantum mechanics/molecular mechanics (QM/MM) simulations were conducted, including the divalent ions, coordinating residues (D839, N863), the catalytic H840, critical DNA bases (G-3, T-4, C-5), and the surrounding waters in the QM region. For each system, ∼20 ps of ab-initio MD at a Density Functional Theory (DFT) level were collected in two replicates. As a result, Ca^2+^ maintains coordination distances similar to those of Mg^2+^, preserving the active site geometry (Fig. 4b, Supplementary Fig. 21). This is consistent with NMR data, where the binding of Ca^2+^ to the HNH-L2 construct results in similar structural perturbations to those observed for Mg^2+^ (Supplementary Fig. 18). Furthermore, the p*K*_a_ of H840 remains largely unchanged in the presence of Mg^2+^ and Ca^2+^ (Fig. 4c, Supplementary Fig. 22), indicating that the effects of Mg^2+^ and Ca^2+^ within the HNH active site are comparable. In contrast, Co^2+^ binding results in a significant reduction in coordination distances, notably displacing H840 from P_SCI_ (Fig. 4b, Supplementary Fig. 21). In the presence of Ca^2+^, the distances involving the ion, the P_SCI_ and the water nucleophile (WAT_nu_) – Ca^2+^–P_SCI_, P_SCI_–WAT_nu_, and WAT_nu_–H840 – are consistent with the reference Mg^2+^-bound system, preserving the active site’s structural integrity. Contrarily, with Co^2+^, shortened Co^2+^–P_SCI_ and P_SCI_–WAT_nu_ distances are accompanied by an increased WAT_nu_–H840 distance. This misalignment disrupts the catalytic geometry, potentially lowering the nucleophilicity of WAT_nu_ and diminishing its ability to facilitate DNA cleavage.

**Figure 4.**
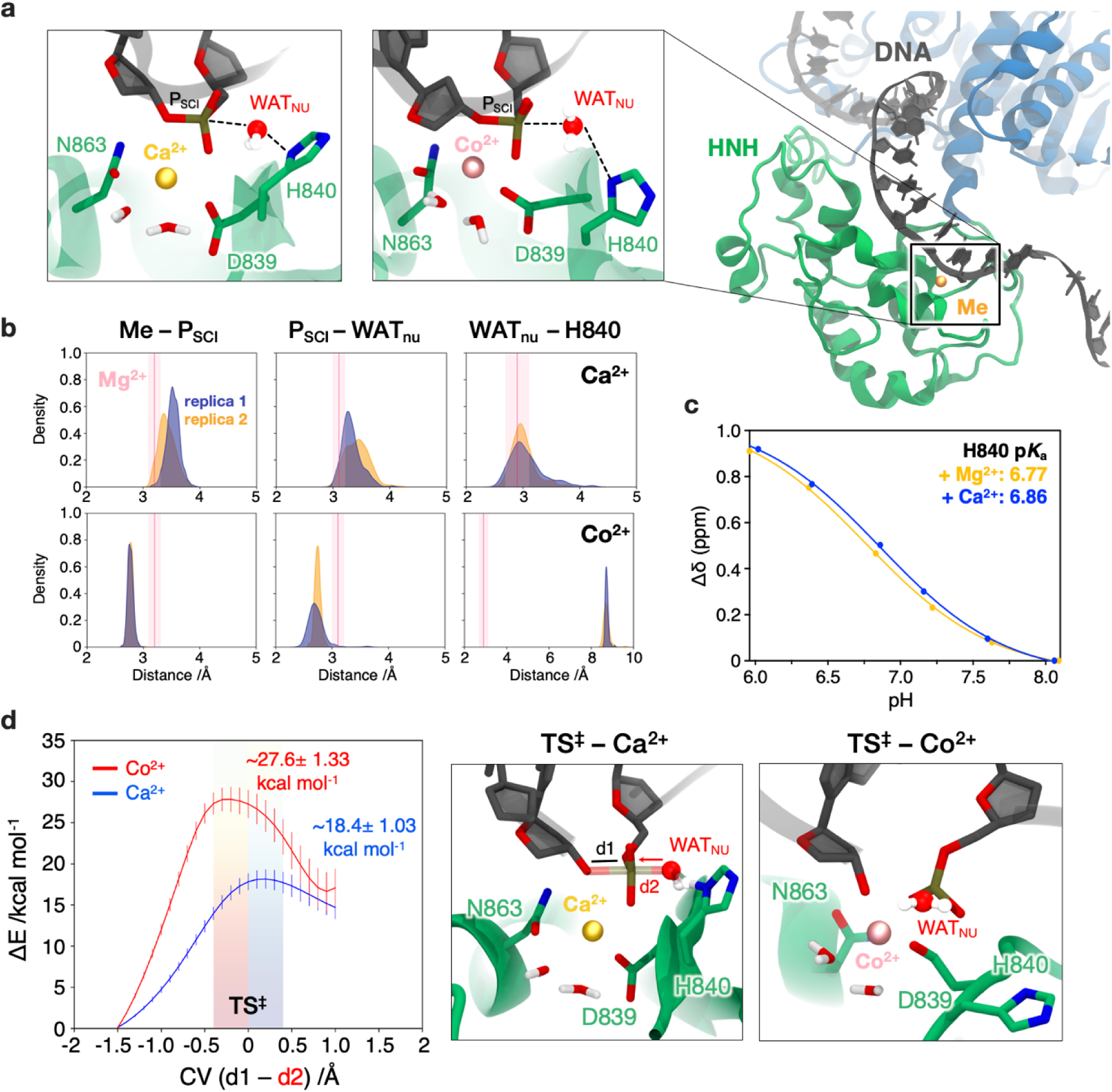
Effect of divalent metal ions on the HNH active site chemistry. **a.** Conformations of the HNH active site in the presence of Ca^2+^ (left) and Co^2+^, obtained from classical molecular dynamics (MD) and ab-initio MD simulations at the QM(DFT-BLYP)/MM level. **b.** Distributions of key distances involving the metal ion (Me), the scissile phosphate (P_SCI_), and the water nucleophile (WAT_nu_), derived from ∼20 ps of ab initio QM/MM MD simulations (two replicates) with Ca^2+^ (top) and Co^2+^ (bottom). The red line in each plot indicates the reference distances obtained in the presence of Mg^2+^, computed using the same methodology; the average values are shown with shaded regions representing the associated error bars. **c.** NMR determined p*K*_a_ of the catalytic H840 in the presence of Mg^2+^ (yellow) and Ca^2+^ (blue). Errors in the chemical shift are smaller than the data points. The p*K*_a_ of H840 could not be quantified when bound to Co^2+^ due to extensive paramagnetic line broadening around the binding site. **d.** Free energy profiles for phosphodiester bond cleavage in the presence of Ca^2+^ (blue) and Co^2+^ (red), obtained through QM(BLYP)/MM MD and thermodynamic integration. The difference in distance between the breaking and forming P–O bonds is used as the reaction coordinate (RC = d1−d2, shown in the right panel). Error bars represent standard deviations derived from error propagation analysis of the dataset, with each data point reflecting the mean over the last ∼6–10 ps of converged ab initio MD. The Transition State (TS**^‡^**) structures in the presence of Ca^2+^ and Co^2+^ are shown on the right.

To assess the impact of these structural differences on catalysis, we conducted free energy simulations, characterizing phosphodiester bond cleavage. A thermodynamic integration approach was combined with QM/MM simulations, using the difference in distance between the breaking and forming P–O bonds as a reaction coordinate. This methodology was previously employed to characterize the catalytic mechanism in the presence of Mg^2+32^.

The free energy profiles indicate that Ca^2+^ supports catalysis with an activation barrier of ∼18.4 ± 1.03 kcal/mol (Fig. 4d, Supplementary Fig. 23), resembling the barrier obtained for Mg^2+^ (∼17.06 ± 1.22 kcal/mol) ^32^. Contrarily, Co^2+^ raises the activation barrier to ∼27.6 ± 1.33 kcal/mol, indicating a less favourable catalytic pathway. This ∼9 kcal/mol higher reaction barrier with Co^2+^ corroborates biochemical results showing an abolition of catalysis in the presence of Co^2+8,18^. A mechanistic analysis elucidates this difference, showing that while Ca^2+^ follows an associative S_N_2 mechanism (similar to Mg^2+^), Co^2+^ drives the reaction towards a dissociative mechanism due to the misalignment of H840. Indeed, in the presence of Ca^2+^, H840 effectively abstracts a proton from WAT_nu_, facilitating the nucleophilic attack on P_SCI_ with simultaneous breaking of the O3’—P bond and forming the P—O_WAT_ bond (Fig. 4d, Supplementary Figs. 23±-25). Conversely, in the presence of Co^2+^, a premature O3’—P bond cleavage is followed by a proton transfer from WAT_nu_ to O3’, resulting in a less efficient reaction pathway. To further validate the catalytic mechanism with Ca^2+^, metadynamics simulations were performed (Supplementary Fig. 26), describing phosphodiester bond cleavage and deprotonation of the nucleophilic water across two dimensions. The free energy surface confirmed an S_N_2-like mechanism and an activation barrier of ∼18.5 ± 0.91 kcal/mol, similar to that computed for Mg^2+^ using the same approach (∼17.4 ± 0.84 kcal/mol)^32^, underscoring that divalent metals retaining the integrity of the active site preserve the catalysis.

## Discussion

Here, molecular simulations and biochemical assays were used to elucidate how divalent metal ions influence conformational activation and target DNA cleavage in CRISPR-Cas9. Free energy simulations and Markov State Models delineate the thermodynamic and kinetic mechanisms driving activation of the catalytic HNH domain under varying concentrations of Mg^2+^ ions. We show that at high Mg^2+^ levels (10 mM), HNH transitions more readily from the pre-active to active conformation with the rate-limiting step occurring nearly twice as fast compared to low Mg^2+^ conditions (10 μM) (Figs. 1-2). This aligns well with the ∼2.5-fold difference in transition rates observed in FRET experiments^18^, and is consistent with kinetic studies^17^, indicating that Cas9 fails to reach the active state without Mg^2+^. Single-molecule analyses have also shown that HNH adopts distinct conformations depending on the presence of Mg^2+19^, and rarely transitions to the active state in the absence of divalent cations^18^.

Extensive MD simulations uncover the formation of a divalent metal binding pocket (DBP) at the HNH-RuvC interface, involving residues from the L2 region, the RuvC domain, and the DNA NTS (Fig. 3). Such “cryptic pockets” – transient cavities not apparent in static crystal or cryo-EM structures – can emerge from MD simulations, shedding light on their functional significance^25–27^. Here, this cryptic DBP forms at high Mg^2+^ concentration and is pivotal in regulating HNH dynamics. Indeed, elevated Mg^2+^ levels increase ionic residence time at this site, facilitating the conformational changes necessary for HNH activation. Similar effects are observed with Ca^2+^ and Co^2+^ (Supplementary Fig. 15), where high concentrations likewise drive the structural rearrangements necessary for activation. These findings highlight a general mechanism by which divalent cations induce conformational activation through cryptic site stabilization.

The cryptic DBP critically involves the L2 region, which, along with L1, forms flexible linkers tethering HNH to RuvC^13^. These linkers, absent in early experimental structures due to their inherent flexibility^3^, are observed at higher resolution in cryo-EM structures^14,15^ and act as signal transducers in the coupled catalytic action of HNH and RuvC^13,23,29,31^. Our simulations show that at high Mg^2+^ levels, structural rearrangements of L2 facilitate HNH activation through bridging interactions at the DBP, formed by Mg^2+^, L2, RuvC, and the NTS (Fig. 3). This reveals that divalent metals not only promote HNH activation but are key to mediate critical communication between HNH and RuvC. Mutation of the DBP residues markedly reduce the cleavage activity of both HNH and RuvC, and consequently the intermediate nicked DNA species, indicating that alterations of the DBP disrupt the delicate balance necessary for the coupled catalytic action of HNH and RuvC^13,23,29–31^. Moreover, mutations at distinct DBP sites (i.e., M1 and M3 in RuvC, M2 in the L2 region of HNH) consistently reduce RuvC cleavage activity and reveal strong correlation between the formation of linear DNA species and diminished HNH catalysis (Fig. 3h). This suggests that such mutations impair HNH dynamics and hinder the HNH-RuvC communication. Divalent metals thereby stabilize interactions within the DBP, enhancing coordination between HNH and RuvC, and facilitating the HNH domain activation.

Studies of the effects of Mg^2+^, Ca^2+^, and Co^2+^ of TS cleavage using quantum-classical simulations reveal distinct differences. Mg^2+^, the optimal catalytic ion, coordinates with key residues and waters to promote DNA cleavage via an associative S_N_2 mechanism (Fig. 4). Ca^2+^ maintains this coordination geometry and catalyses TS cleavage through a comparable mechanism. Contrarily, Co^2+^ induces significant structural deviations, shifting the reaction toward a dissociative mechanism and increasing the activation barrier, ultimately impairing catalysis. This demonstrates that the catalytic activity is metal-dependent, while the ionic strength, independent of the specific ion, promotes the conformational activation of Cas9.

In summary, this study provides key insights into how divalent metal ions regulate the conformational activation and target DNA cleavage in CRISPR-Cas9. By integrating free energy simulations, Markov State Models, and quantum-classical approaches, biochemical and NMR experiments, we show that divalent metals such as Mg^2+^, Ca^2+^, and Co^2+^ promote the conformational activation of the catalytic HNH domain. These ions bind within a cryptic divalent metal-binding pocket (DBP), which emerges dynamically in simulations and plays a central role in stabilizing HNH conformations and coordinating its activity with RuvC. The DBP thus serves as a critical regulatory hub, mediating metal-dependent communication between the two nuclease domains. These findings uncover an essential regulatory mechanism in Cas9 function and offer key insights into the Cas9-mediated genome editing in cells and organisms where ion availability and metal homeostasis vary widely.

## Methods

### Molecular Dynamics simulations

Molecular dynamics (MD) simulations were performed on two distinct structural states of CRISPR-Cas9: the pre-active conformation (PDB 5F9R, solved at 3.40 Å resolution^13^) and the catalytically active conformation (PDB 6O0Y, solved at 3.37 Å resolution^14^), representing two distinct states of the HNH domain. Each system was fully solvated in explicit water and neutralized with divalent metal ions (Mg^2+^, Ca^2+^, or Co^2+^). To model varying ionic environments, simulations included either 13 MCl_2_ units (M = Mg^2+^, Ca^2+^, or Co^2+^) corresponding to a high ion concentration of 10 mM, or a single MCl_2_ unit to represent a low concentration of 10 μM. The resulting systems comprised ∼340,000 atoms within simulation boxes of ∼180*120*140 Å^3^. MD simulations were preformed using a protocol tailored for protein/nucleic acid complexes^33^ and applied in studies of CRISPR-Cas systems^34–37^. The Amber ff19SB^38^ force field was employed, incorporating the OL15^39^ and OL3^40^ corrections for DNA and RNA, respectively. The TIP3P^41^ model was used for water molecules. The Li & Merz model was used for ions^42,43^, describing the metal sites in agreement with QM/MM simulations (Supplementary Fig. 21). The 12-6-4 polarizable Li-Merz^44,45^ and the Allnér^46^ parameters were also used to model Mg^2+^ ions in the pre-active state. An integration time step of 2 fs was used. All systems were minimized and equilibrated through a staged heating protocol, gradually raising the temperature to 300 K under positional restraints that were progressively released. Production runs were conducted in the NVT ensemble, with three ∼1 μs replicates for each condition, resulting in ∼48 μs of simulation time. The GPU-empowered version of AMBER 22^47^ was used as MD engine. Full details are reported in the Supplementary Methods.

### Umbrella Sampling simulations

The Umbrella sampling (US) free energy method was employed to characterize the conformational transition of the HNH domain from the pre-active to active state^22^. In this approach, a number of simulations (US windows) are run in parallel with additional harmonic bias potential applied to selected Reaction Coordinates (RCs):

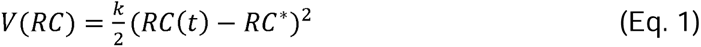

where *V*(*RC*) is the value of the bias potential, *k* is a bias force constant, *RC*(*t*) is the value of *RC* at given time *t* and *RC*\* is the reference value of *RC*. By using different *RC*\* values in each US window, one can sample the biased probability distribution *p_b_*(*RC*) along the whole *RC* range of interest. The difference in root-mean-square deviation (RMSD) of the Cα atoms in HNH and portions of REC (residues 175-393, 778-906) relative to the pre-activated and active states, was used as a RC (Supplementary Fig. 2a). Two independent US simulations were performed: under high (10 mM) and low (10 μm) concentrations of Mg^2+^. The systems were simulated in 36 overlapping windows, with the RC ranging from 7.4 Å to 0.2 Å, using a harmonic restraint with a spring constant of 50 kcal/mol·Å^2^. The centre of the harmonic bias potential was distributed along the RC in 36 windows, each separated by 0.2 Å, ensuring proper overlap of the probability distributions. For each system, ∼120 ns-long trajectories were obtained per US window, resulting in ∼4.3 µs of collective sampling per system (and a total of ∼8.6 µs). The free energy profiles were computed using the Weighted Histogram Analysis (WHAM) method^48^, discarding the first ∼20 ns of each window to ensure convergence. Error estimation on the free energy profile was performed using Monte Carlo bootstrap error analysis^49^. Details are reported in the Supplementary Methods.

### Markov State Models

To resolve the kinetics of the HNH transition, Markov State Models (MSMs)^20,21^ were constructed using unbiased MD simulations initiated from 36 representative conformations extracted from the US trajectories. Each seeded trajectory was propagated for ∼100 ns, totalling ∼3.6 μs per condition. Features used for MSM construction included inter-residue Cα distances (S355–S867 and S867–N1054), previously validated by FRET studies. Time-lagged Independent Component Analysis (TICA)^24^ was performed (lag time = 5 ns), followed by k-means clustering into 100 microstates^50,51^, selected based on the saturation of the VAMP-2 score^52^. Implied timescales analysis^53^ confirmed a Markovian lag time of 5 ns, which was used to construct Bayesian MSMs. The Chapman–Kolmogorov test validated the models’ Markovianity^54^. Macrostates were defined using PCCA++ clustering^55^, identifying four metastable states. MSM-derived stationary distributions were used to reweight MD configurations, enabling the calculation of MSM-reweighted free energy landscapes projected onto the TICA space. Finally, Transition Path Theory (TPT)^56,57^ was applied to estimate the dominant flux pathways between the pre-active and active states based on forward and backward committor probabilities. MSM analysis was performed through the PyEMMA package^58^.

### QM/MM simulations

QM/MM simulations were performed on the active conformation of the HNH domain in the presence of Ca^2+^ and Co^2+^ in the active site. The QM region comprised the divalent ion, coordinating residues (D839, N863), the catalytic H840, DNA bases G-3, T-4, C-5, and nine water molecules, including the nucleophile and ion-coordinating waters. This region totalled 123 atoms and was capped with hydrogen atoms to saturate valence at the QM–MM boundary. H840 was modelled in its neutral ε-tautomer, consistent with prior QM/MM simulations including Mg^2+ 32^. QM atoms were treated at the DFT/BLYP^59,60^ level; the MM region used the same force field as in classical MD simulations (*vide supra*). Ab-initio MD simulations were carried out using the CPMD version 4 code^61^. The wave functions were expanded in a plane wave basis set up to a cutoff of 75 Ry. Interactions between the valence electrons and ionic cores were described using norm-conserving Martins-Troullier pseudopotentials^62^. Electrostatic coupling between periodic QM images was removed using Tuckerman’s scheme^63^. A Hamiltonian treatment of the electrostatic interaction between the QM and MM regions was used^64^. The QM/MM protocol included initial wavefunction optimization, followed by ∼6 ps of Born–Oppenheimer MD at 300 K in NVT ensemble, using a 20 au (∼0.48 fs) time step. This was followed by Car–Parrinello MD using a 5 au (∼0.12 fs) timestep and a fictitious electron mass of 600 au. For each system, ∼20 ps of unconstrained QM/MM simulations were performed. The active site conformations derived from these simulations were subsequently used as starting points for free energy simulations to study the catalytic mechanism (*vide infra*).

### Thermodynamic Integration

Thermodynamic Integration and the “blue moon ensemble” method ^65,66^ were used to compute the free-energy profiles for phosphodiester bond cleavage in the active state of HNH in the presence of Ca^2+^ and Co^2+^. Here, the average converged constraint forces are computed and integrated along a given RC, deriving the associated free energy profile. The difference in distance between the breaking and forming P–O bonds was used as RC, in line with studies of DNA cleavage by HNH in the presence of Mg^2+32^. Starting from a RC = −1.5 Å, corresponding to the reactant state, a total of 26 sequential windows were sampled along the RC at 0.1 Å intervals, extending to the product state (RC = 1.5 Å). Each window was simulated for ≥10 ps, ensuring convergence of the constraint force (Supplementary Fig. 24), and resulting in a total of ∼260 ps of ab initio MD. To estimate the error associated to hysteresis, we also computed the backward free energy profiles. These simulations were initiated from the post-transition state region (RC = 0.5 Å) and involved sampling 8 windows per system in the reverse direction along the RC. These simulations resulted in additional ∼80 ps of sampling for each system. The statistical error at each point along the forward and backward free energy profiles was estimated using error propagation analysis. The total uncertainty in the free energy barrier was then obtained by combining the statistical error with the hysteresis error, calculated as the discrepancy between the forward and backward pathways. Details are in the Supplementary Methods.

### Metadynamics

To further elucidate the catalytic mechanism in the presence of Ca^2+^, QM/MM metadynamics^67^ simulations were performed on the Ca^2+^-bound active HNH complex. Here, a time-dependent biasing potential is added to the system’s Hamiltonian, applied as a function of selected collective variables (CVs) that represent the process of interest. Two CVs were employed: (i) the difference in distances between the breaking and forming P–O bonds, and (ii) the distance between the δ-nitrogen of H840 and the hydrogen atom of the nucleophilic water molecule. An extended Lagrangian formulation of the method was employed to ensure consistent coupling with the QM/MM framework^68^. For both collective variables, the fictitious particle mass was set to 16 amu, and the associated force constant was 0.24 au. The height of the Gaussian terms was 0.5 kcal/mol, while their width was set at 0.05 Å, consistent with the oscillations of the CVs observed in unbiased Car-Parrinello QM/MM simulation. A total of ∼5000 Gaussians were deposited during the simulation, yielding a total of ∼120 ps of metadynamics sampling. Multiple transitions between reactant and product states were observed (Supplementary Fig. 26), confirming thorough sampling. Free energy profiles were computed by averaging over ∼10 ps blocks; errors were estimated using block averaging from the converged region. The statistical error associated with the free energy for phosphodiester bond cleavage was estimated using average blocking over ∼10 ps segments from the converged portion of the metadynamics trajectory. Details are reported in the Supplementary Methods.

### In Vitro Cleavage

Mutants M1 (D54A/S55A/E57A), M2 (Q920A/E923A), and M3 (T13A/N14A) were generated by site-directed mutagenesis of the SpCas9 plasmid (Addgene pMJ806) using primers listed in Supplementary Table 1. Recombinant wild-type and mutant Cas9 proteins were expressed in *E. coli* BL21(DE3) cells, induced with 0.5 mM IPTG for 18 hr at 18 °C, and purified by Ni-affinity chromatography followed by TEV tag cleavage and SP cation exchange chromatography. Proteins were concentrated, aliquoted, flash-frozen, and stored at −80 °C. Cas9 was pre-incubated with gRNA in 1.5X excess in cleavage buffer [20 mM Tris-HCl pH 7.5, 100 mM KCl, 5% glycerol, 1 mM DTT, 10 mM MgCl_2_] for 10 min at room temperature. Plasmid cleavage reactions were initiated at 37 °C combining 100nM Cas9-gRNA with 2 nM circular plasmid containing an NGG PAM and Lambda1 target sequence. Aliquots were quenched at defined time points in 0.5 M EDTA and resolved on 1% agarose gels. Band intensities for supercoiled (SC), nicked (Ni), and linear (Li) species were quantified in FIJI and fit by non-linear regression using KinTek software: SC to a single exponential, Ni and Li to a double exponential. To generate a DNA substrate containing the NGG PAM and Lambda1 target sequence, fluorescently labelled oligos (5′-FAM label on TS or NTS) were annealed with their complement. Cas9–gRNA complexes were assembled as above and oligoduplex cleavage reactions initiated with 10 nM duplex at 37 °C. Aliquots were quenched in EDTA and analysed by capillary electrophoresis (ABI 3130xl) to quantify fraction cleaved over time. Data were fit to a single exponential using GraphPad Prism to determine *k*_obs_ for TS and NTS cleavage. The sgRNA sequence used in cleavage experiments was 5′-GACGCAUAAAGAUGAGACGC + 80-mer SpCas9 scaffold-3′. The dsDNA substrates were prepared from complementary TS (5′-AGCTGACGTTTGTACTCCAGCGTCTCATCTTTATGCGTCAGCAGAGATTTCTGCT-3′) and NTS (5′-AGCAGAAATCTCTGCTGACGCATAAAGATGAGACGCTGGAGTACAAACGTCAGCT-3′) oligonucleotides. For fluorescence detection, either the target strand or the non-target strand carried a 5′-FAM label, depending on whether target strand or non-target strand cleavage was being resolved.

### NMR Spectroscopy

The HNH-L2 domain (residues 772–925) of *S. pyogenes* Cas9 was cloned into pET-28a(+) with an N-terminal His₆-tag and TEV cleavage site. Protein was expressed in M9 minimal medium supplemented with ^15^NH_4_Cl and ^13^C-glucose, induced with 1 mM IPTG, and purified using Ni-NTA affinity and size-exclusion chromatography (full details in the Supplementary Information). Post-purification, protein was dialyzed into buffers containing KCl, MgCl_2_, or CaCl_2_ depending on the sample condition. NMR experiments were conducted at 600 MHz and 25°C using a Bruker Avance NEO spectrometer. Backbone assignments were obtained using standard triple-resonance experiments (full details in Supplementary Methods)^69^ and validated against BMRB entry 27949^70^. Chemical shift perturbations were computed from ^1^H–^15^N HSQC spectra^71^ and parameters from *T*_1_, *T*_2_, ^1^H-[^15^N]-NOE, and CPMG relaxation dispersion experiments were measured using established protocols^72,73^ with uncertainties were derived from replicate spectra^74–76^. Histidine p*K*_a_ values were determined from ^1^H–^13^C HMQC titrations and fitted using a modified Henderson–Hasselbalch equation, as previously described^32,77^.

## Acknowledgments

This material is based upon work supported by the NIH (Grant No. R01GM141329, to GP; Grant No. R01GM136815, to GP and GPL; and Grant No. R35GM138348 to DWT) and the NSF (Grant No. CHE-2144823, to GP and Grant No. MCB-2143760, to GPL). GP acknowledges support by the Alfred P. Sloan Foundation (Grant No. FG-2023-20431) and the Camille and Henry Dreyfus Foundation (Grant No. TC-24-063). ES was supported by a Blavatnik Family Graduate Fellowship. The computational studies performed here were carried out using Expanse at the San Diego Supercomputing Centre through allocation MCB160059 and Bridges2 at the Pittsburgh Supercomputer Centre through allocation BIO230007 from the Advanced Cyberinfrastructure Coordination Ecosystem: Services & Support (ACCESS) program, which is supported by NSF support grants #2138259, #2138286, #2138307, #2137603, and #2138296.

## Author Contribution

MA performed molecular simulations, analysed the data, and wrote the manuscript. AS analysed the data. DS performed and analysed biochemical experiments, aided by IS. ES carried out NMR sample preparation and experiments, analysed the data, and contributed to manuscript writing. GPL supervised the NMR experiments. DWT supervised the biochemical experiments. GP supervised the computational studies, conceived the research, and contributed to manuscript writing. All authors critically reviewed and edited the manuscript.

## Competing Interests

The authors declare no competing interests.

## Data Availability Statement

Atomic coordinates of the optimized computational models are available in figshare with the identifier https://doi.org/10.6084/m9.figshare.19158080. NMR resonance assignments for the HNH nuclease are available in the BMRB entry 27949. All other data are available from the authors upon request.

## Notes

### Competing Interest Statement

The authors have declared no competing interest.

### Summary of Updates

This version includes an additional author, Alexa L. Knight, who is contributing NMR and biochemical experiments.

